# Intraspecific microbiome dynamics across the life cycle of the milkweed bug *Oncopeltus fasciatus*

**DOI:** 10.1101/2024.10.28.620616

**Authors:** Will Larner, Nádia Thölke da Silva Grego, Kristen A. Panfilio

## Abstract

The microbiome is an important part of the complete nutritional and genomic profile of insects. The species-rich insect order Hemiptera (aphids, cicadas, true bugs) is highly diverse for mode of microbiome acquisition, with the conundrum that species in the seed-feeding subfamily Lygaeinae have lost obvious anatomy for housing bacteria, either in bacteriocytes or midgut crypts. Here we characterize the microbiome of the milkweed bug *Oncopeltus fasciatus* as a tractable lygaeinid, using *16S rRNA* sequencing. We assess how bacterial taxa vary between the sexes and across life history stages in a controlled environment, focusing on maternal-to-embryo transmission and distinguishing egg-stage constituents that are superficial or internal (transovarially transmitted). Among a core microbiome of 28 genera, the egg stage shows the greatest diversity, with a particular expansion of the family Comamonadaceae. We also analyze inter-individual variability in nymphs and adults and validate structured, stage-specific detection of seed material. Comparative analysis identifies Rhizobium as a notable microbiome constituent in seed-feeding Hemiptera, which we had previously shown to lack nitrogen metabolism components in the genome. Overall, we provide a nuanced assessment of bacterial abundance dynamics between individuals and across the life cycle and discuss the implications for acquisition and potential relevance as nutritional endosymbionts. This will underpin comparative investigations in seed-feeding bugs and future work in *O. fasciatus* on tissue-specific and diet-specific microbiome profiles, including in natural populations.

## INTRODUCTION

Insects are integral components of food webs and ecosystems across the globe [1–3]. With insect species’ abundances in flux as their geographical ranges change, in part linked to changes in local climate and human land use [4, 5], understanding the dynamics of biodiversity has become a key challenge. One avenue to investigate this is to elucidate the genomic basis of insect feeding ecology, in terms of species’ capacities to exploit diverse or novel food sources or distinct ecological niches [6–9].

With >250 sequenced insect genomes to date [10], comparative genomics can reveal molecular evolutionary changes associated with ecology. This includes expansions, losses, or rapid evolution in protein families with roles in environmental sensing (chemoreceptors, opsins), detoxification (antioxidants, antimicrobial peptides), and metabolism (digestive enzymes) [11–13]. For example, gene repertoire changes have been linked to dietary shifts between species [7] and between whole suborders of insects [14]. Mutations in specific metabolic enzymes underpin convergent adaptation by different species to the same host plant [e.g., 15, 16]. Importantly, new genes acquired by lateral gene transfer ([11, 17, 18]) or acquired directly by insects’ microbial endosymbionts [19, 20] are also associated with changes in food source and geographic range. In contrast, the loss of gut microbiota correlates with impaired growth and fitness [21, 22]. Thus, meaningful genomic profiling should encompass not only insects’ own protein-coding repertoires, but also the extended genomic profile of the insect holobiont, including the microbiome [23–25].

Integrated genomic profiling is particularly important for understanding the complex molecular basis of insect phytophagy, or plant feeding, strategies [14, 19]. Phytophagy is widespread but also presents nutritional challenges. Food quality may be poor and lack essential vitamins or amino acids [20]. Plant tissues are protected by tough cell walls [17] and may contain toxic compounds such as cardenolides or other cardiac glycosides [15, 16].

The Hemiptera is the most species-rich order of hemimetabolous insects, with members including aphids, psyllids, and true bugs (Heteroptera) [26]. Secondary reacquisition of phytophagy in the true bug infraorder Pentatomomorpha (Fig. 1, [26]) has led to radiations of species that are invasive and polyphagous [18, 27], preferentially specialist [28], or strictly monophagous [29] for plant species and tissue. How do feeding ecology types compare for the metabolic repertoires of the insect itself or its microbial constituents?

**Figure 1.**
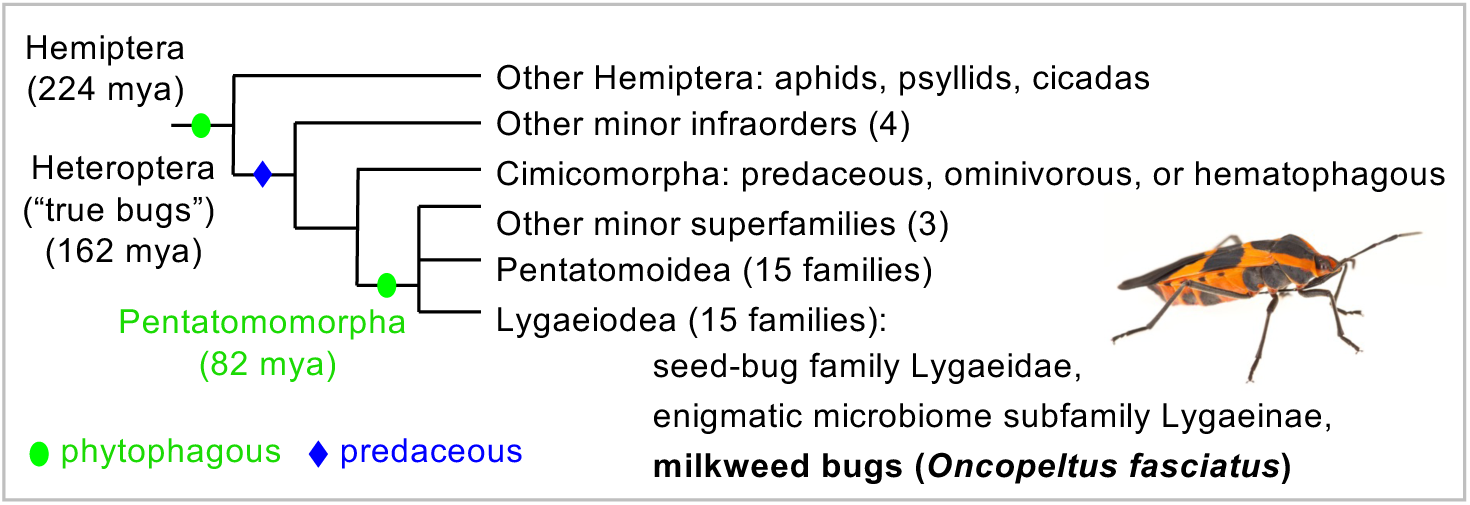
Feeding ecology evolutionary history and phylogenetic context of *Oncopeltus fasciatus*. For the Lygaeinae, the designation “enigmatic microbiome” refers to the absence of both midgut crypts and of bacteriocytes or bacteriomes as bacteria-housing structures that provide an anatomical framework for inferring microbiome constituents (*i.e.*, extracellular or intracellular, respectively). Taxonomic relationships and feeding strategy changes: [9, 26]. Milkweed bug image: Jena Johnson Photography, as in [11].

Nutritional endosymbionts have been characterized in many Hemiptera [reviewed in 30, 31], including prominent examples such as aphids, with a nutrient-poor, fluid-feeding lifestyle [20, 32]. Particularly within the true bug infraorder Pentatomomorpha, there is extensive diversity for endosymbiont transmission type, dedicated host anatomy, and microbe taxonomy [29, 33]. Briefly, bacterial acquisition is often environmental, through nymphal feeding at each generation [34], primarily via ingestion of feces (coprophagy) from conspecific individuals [35]. Another predominant route of symbiont transmission is via hatchling ingestion of bacteria vertically provided by the mother, such as through egg smearing or capsule or jelly secretions external to the eggs [36, 37]. Vertical transmission can also be transovarial, with bacteria deposited by the female directly into the egg [38]. Within the insect’s abdomen, bacteria may be harbored in dedicated midgut crypts, which is typical for extracellular *Gammaproteobacteria* in Pentatomoidea [39]. Alternatively, intracellular *Burkholderia* symbionts in Lygaeoidea are housed in bacteriocytes that are often distinctively pigmented and located near the gonads [40].

The seed-feeding subfamily Lygaeinae are a notable exception, as specialist and monophagous feeders with an enigmatic, under-studied microbiome (Fig. 1). This subfamily lost midgut crypts, but reports differ for bacteriocyte presence, even within a genus [29, 33]. Older sequencing analyses suggest these insects have a diverse microbiome [29, 33, 39], and this awaits full characterization. The milkweed bug *Oncopeltus fasciatus* is a specialist feeder and research model for physiology, development, and evolutionary ecology since the mid twentieth century [reviewed in 11, 28, 39]. Adult anatomy and embryology are well described, corroborating the absence of obvious microbiome-housing tissues [41–44]. Furthermore, genome analysis identified an extensive metabolic enzyme repertoire, but with some notable omissions, such as in nitrogen metabolism and amino acid synthesis [11].

What is the composition of the *O. fasciatus* microbiome and what proportions of the full complement derive from distinct acquisition strategies at different life history stages? Here we characterize the microbiome profile across the life cycle, using *16S rRNA* amplicon sequencing. Our sampling focused particularly on maternal and embryonic stages, and we distinguish microbial constituents within the egg or on the eggshell surface. Furthermore, to control for potentially distinct ecological microniches between embryonic, nymphal, and adult stages [28], and for precise assessment of inter-individual variability [45], we assayed bugs from a uniform laboratory colony environment. We hypothesized that the *O. fasciatus* microbiome is primarily acquired via transovarial transmission, as in other Lygaeoidea [29, 33], slightly augmented through postembryonic environmental acquisition [30, 46]. In fact, we find high inter-individual nymphal variation and impoverished but sex-specific adult profiles, contrasting with diverse egg microbiomes with both transovarial and external taxa.

## MATERIALS AND METHODS

### Milkweed bug culture and life history samples

A laboratory colony of *Oncopeltus fasciatus*, cultured continuously since June 2014, was used (Carolina Biological Supply strain, Burlington, North Carolina, USA, [as used in, e.g., 8, 11, 47]). Generational cohorts of bugs were maintained in plastic cages at 25 °C, with a 12:12 h light:dark cycle, and provisioned with water, peeled organic sunflower seeds (Holland and Barrett, Nuneaton, UK), and loose cotton wool for oviposition (Robinson Healthcare, Worksop, UK). Material for all biological replicates of a given sample type was collected on the same day from the same cohort.

Seven life history stages and egg treatments were analyzed (Fig. 2A). Embryonic samples were collected within the first day (young eggs, EY, 0-24 h) or the second half of embryogenesis (old eggs, EO, 3-7 d; precisely: 70-165 h), and assayed as untreated eggs that retained external bacterial constituents (EY, EO) or after surface sterilization (washed eggs: EYW and EOW) to retain only egg-internal constituents. Nymphs (N) were assayed at the final (fifth) juvenile instar [48], as in previous true bug work [39]. Within this cohort (23-29 d), younger nymphs were selected by size for consistency (<8 mm body length, Fig. 2B). Reproductively active adults (2-4 weeks after the final molt) were sexed by abdominal morphology and pigmentation [41, 49] for the male (M) and female (F) samples.

**Figure 2.**
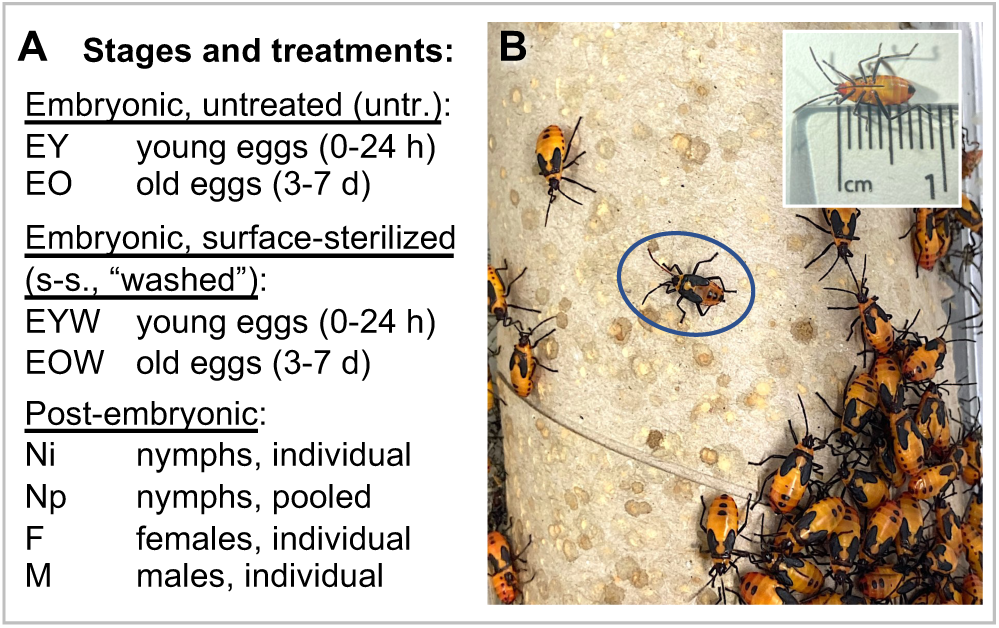
Life history stages and treatment conditions for microbiome profiling. **(A)** All treatment conditions are listed, including the ages of the embryonic samples and which postembryonic samples were based on single individuals or pooled samples (three bugs were used in the pooled nymph samples). Analyzed nymphs were from early in the fifth (final) juvenile instar. **(B)** Images of fifth-instar nymphs. Nymphs used in this analysis were approximately 8 mm in length (inset) and thinner than late-instar nymphs close to the adult molt (compare the circled early-instar nymph with others in the cohort). The bugs congregate on a cardboard tube provided for spatial structure within the cage. The light and dark brown marks on the cardboard are from the bugs’ feces, a standard feature of the colony environment.

### Tissue preparation and DNA extraction

For embryonic stages, to avoid contact with the surface of the eggs, gloves were worn while extracting eggs from the cotton using a fine paintbrush and lightweight forceps. Eggs for the different biological replicates were evenly distributed among sterile 1.5-ml tubes, and dry weight was determined. Surface-sterilization followed a modified protocol [after 39, 40, 50, 51]: eggs were washed in ethanol (for >1 minute each: once in 100% ethanol and twice in 70% ethanol/dH_2_O) followed by three 2-minute rinses in dH_2_O.

Nymphs and adults were anesthetized using CO_2_ and dissected under tap water in a sterile glass dish using sharp forceps. For nymphs, the legs were removed, and tissue was extracted by squeezing the posterior abdomen. For adults, the legs, wings, and abdominal exoskeleton were removed, ensuring intact gut material was collected. Tissue weight was determined in minimal residual liquid (range: 13.5-33.2 mg per insect). Adult samples were processed with one individual per biological replicate. For the smaller nymphs, replicates of either individual nymphs (Ni) or pooled samples of three nymphs (Np) were processed.

Additionally, environmental sampling was performed on laboratory food stocks. The sunflower seeds, crushed in a mortar and pestle, and milled wheat flour (Wright’s Bounty Premium Bakers White Bread Flour, Wright and Sons, Harlow, UK) were weighed to an input of 25 mg.

All samples were processed using the Quick-DNA/RNA Miniprep kit (Zymo Research, Orange, California, USA). DNA quality and quantity were confirmed by spectrophotometry (Implen NanoPhotometer N60/N50, Munich, Germany).

### *16S rRNA* amplicon sequencing and taxonomic classification

As in prior insect microbiome characterization [51], an established Illumina sequencing pipeline was followed (“16S Metagenomic Sequencing Library Preparation”, version 15044223 Rev. B). Briefly, an approximately 425-bp portion of the *16S rRNA* gene was amplified with slightly degenerate primers targeting the V3 and V4 hypervariable regions (forward primer: 5′-tcgtcggcagcgtcagatgtgtataagagacagCCTACGGGNGGCWGCAG-3′, reverse primer: 5′-gtctcgtgggctcggagatgtgtataagagacagGACTACHVGGGTATCTAATCC-3′; adapter sequence in lowercase, 16S-specific sequence in uppercase). PCR was performed according to the protocol, modified to include technical replicates with REDTaq Ready Mix (Sigma/Merck, Darmstadt, Germany) or KAPA Taq EXtra HotStart ReadyMix PCR Kit (Kapa Biosystems, Wilmington, Massachusetts, USA). Agencourt AMPure XP PCR purification with magnetic beads (Beckman Coulter, Brea, California, USA) was used.

For 3-5 biological replicates per sample (2 technical replicates each), sequencing was conducted with an Illumina MiSeq machine, using the v.3 kit with 2x 300-bp reads. Reads were taxonomically classified with Illumina’s 16S Metagenomics app v.1.1.0, through the BaseSpace implementation of the Ribosomal Database Project TRDPU Classifier ([52]; last accessed 7 October 2022).

A selected dataset was independently Sanger-sequenced. Amplicons were cloned into the pGEM-T Easy vector in JM109 competent cells (Promega, Southampton, UK). For each replicate, 3-5 clones were sequenced (M13 vector primers, forward: 5′-TGTAAAACGACGGCCAGT-3′, reverse: 5′-GGAAACAGCTATGACCATGA-3′). After trimming the 16S primers, clones were classified by blastn at GenBank (last accessed 4 October 2024; see Table S1).

### Statistical data handling and phylogenetic analysis

For the genus diversity profiles and Shannon Species Diversity Index, all bacterial taxa were considered. A threshold of 1% abundance was used for genus presence/absence, taken as the mean of the biological replicates, from the median of the two technical replicates. The threshold for species presence was ≥10 reads in both technical replicates for ≥1 biological sample. Graphical and statistical analyses were conducted in GraphPad Prism (v.10.2.3-10.3.1), including tests for normality and determining applicability of Welch’s correction for parametric pairwise comparisons. Maximum likelihood phylogenetic trees were generated with default settings of the phylogeny.fr analysis pipeline [53].

## RESULTS

### Dataset quality and a general trend of declining microbial diversity across the life cycle

To characterize the milkweed bug microbiome across life history stages (Fig. 2), we first assessed data quality with respect to taxonomic level (Fig. 3A-C). We generated 13.35 Gbp of raw data for 61.1 M reads, with >110,000 classified reads per sample (n=48). On average, >94% of reads were classified to the family or higher level, declining at the genus and species levels (Fig. 3A). Nonetheless, we observed no correlation between read depth and species diversity (Supplementary Data File S1), suggesting sufficient sequencing depth was obtained. Further, the technical replicates were congruent, with only two minor exceptions in genus-based clustering (Fig. 3C: asterisks). Thus, we integrated multiple taxonomic levels to strike a balance between robust read classification and taxonomic resolution.

**Figure 3.**
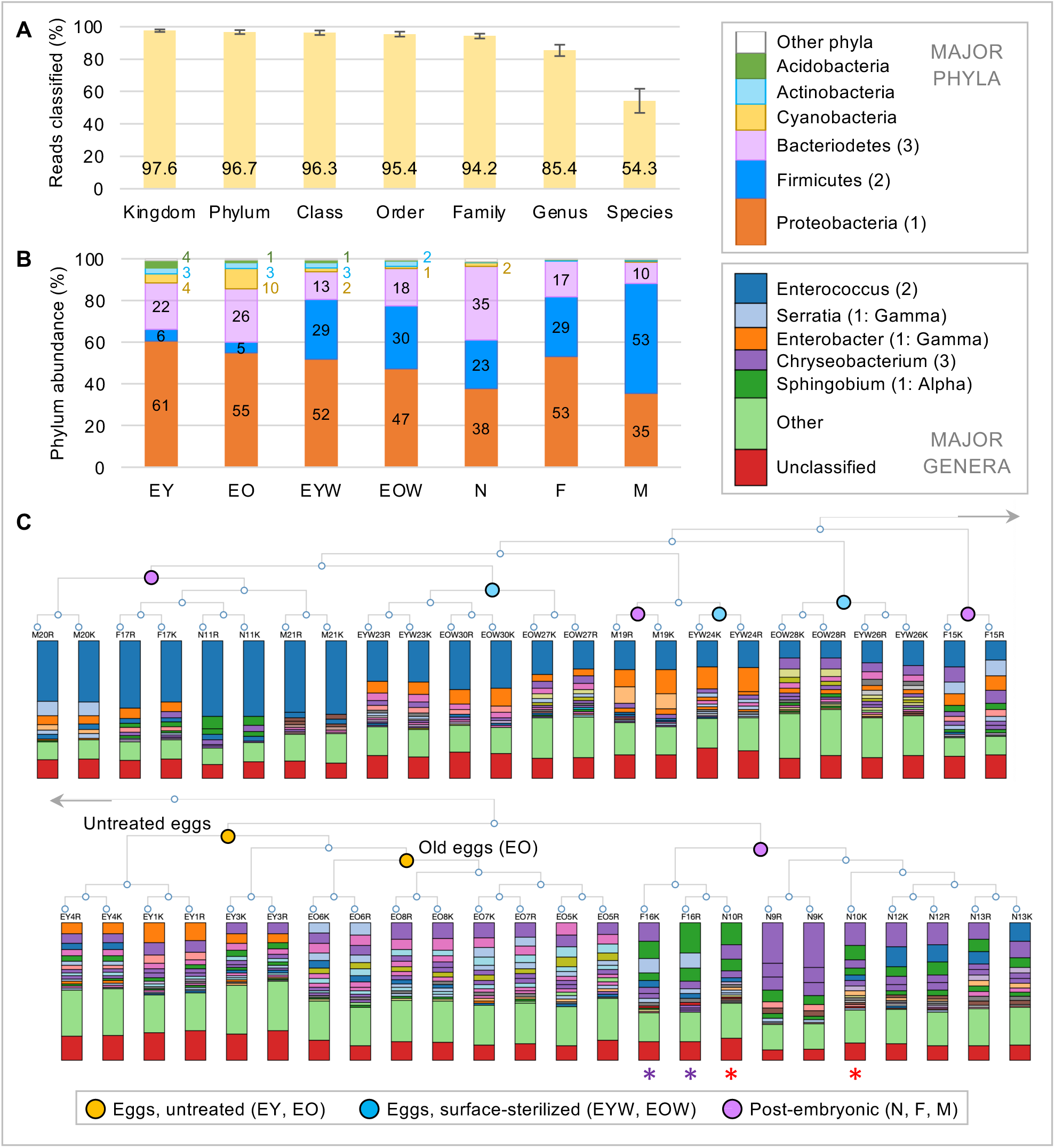
Overview of *16S rRNA* taxonomic classifications. **(A)** Proportion of reads classified per taxonomic level, shown as mean ± standard deviation from all biological and technical replicates (n= 48). The vast majority of reads could be classified, albeit with a noticeable drop-off in classification efficiency at the species level. **(B)** Relative abundance (% reads) of the six most prevalent phyla (upper color legend). **(C)** Genus-level hierarchical clustering dendrogram of all samples, with abundant genera from the top three phyla indicated (lower color legend; “Gamma” and “Alpha” are abbreviations for these classes of Proteobacteria). To accommodate all 48 samples, the dendrogram is split into the two major clusters, which join as indicated by the grey arrows. Colored nodes indicate major clusters for unwashed eggs (orange: with a subcluster for old eggs), surface-sterilized eggs (blue: 3 clusters), and post-embryonic samples (purple: 4 clusters). Asterisks flag the sole instances where technical replicates do not strictly cluster (red: nymphal sample, purple: female sample). Life history treatment abbreviations are as in Fig. 2.

Overall, six bacterial phyla account for >98.7% of the *O. fasciatus* microbiome (Fig. 3B). The Proteobacteria, Firmicutes, and Bacteriodetes are the most abundant (86-99%) and present in all life history samples. These phyla are predominantly represented by just a few major genera (Fig. 3C). Additionally, embryonic – and to a lesser degree nymphal – samples harbored >1% each of the phyla Cyanobacteria, Actinobacteria, and Acidobacteria.

These phylum-level trends are borne out at the genus and species levels for patterns of bacterial diversity and within-sample variation. Untreated eggs that retained surface bacterial constituents, particularly at the older embryonic stage, were the most similar across biological replicates (Fig. 3C) and the most microbially diverse (Fig. 4). Adult samples had significantly lower microbiome diversity (Fig. 4). For genus profile, most nymphal samples clustered with the untreated egg samples (Fig. 3C: lower half of dendrogram). In contrast, all surface-sterilized egg samples were more similar to adult samples, but with no clustering by life history stage (Fig. 3C: upper half of dendrogram). For life history samples derived from a single individual (n=3 per stage), nymphs exhibited greater variability in microbial diversity than either adult males or females (Fig. 4). Overall, this suggests that the egg (surface) is a uniquely diverse microniche, but what is the core microbiome profile, and which bacterial taxa account for major life history stage differences?

**Figure 4.**
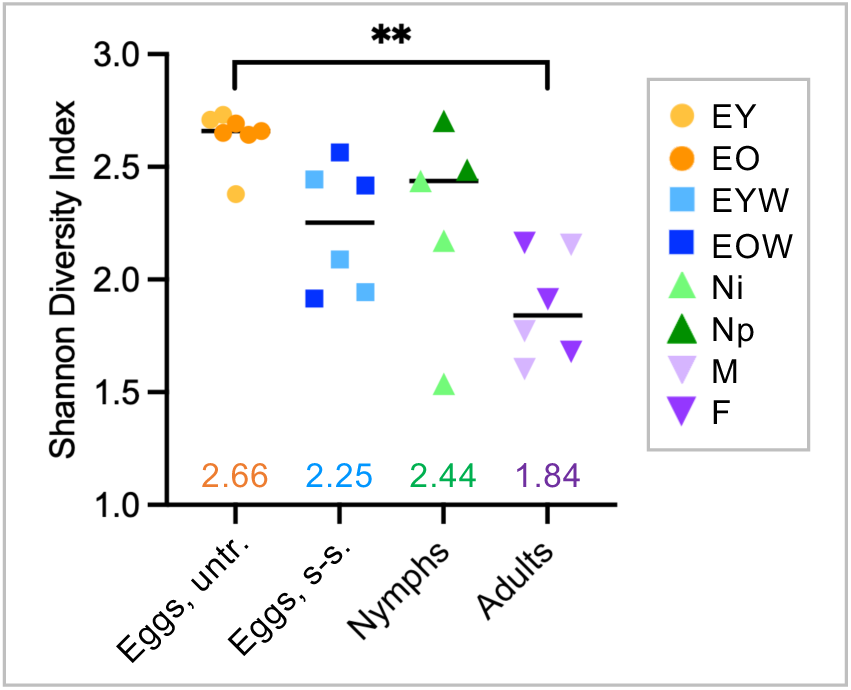
Diversity of the microbial community is greatest in untreated eggs. Shannon Diversity Index of bacterial species composition (accounting for species number and relative abundance) plotted for each biological replicate of the four major life history treatments analyzed. Median values are indicated by the horizontal black bars and specified in colored text. Light and dark plot points distinguish subcategories, as indicated in the legend (abbreviations as in Fig. 2). Significance determined by Kruskal-Wallis and Dunn’s test, with the sole significant difference between untreated eggs and adults (alpha = 0.05, *p_adj_* = 0.0021).

### The core milkweed bug microbiome is phylogenetically diverse

To define a core microbiome profile, we considered all prokaryotic genera with >1% abundance in at least one of the four major life history sample types (untreated eggs, surface-sterilized eggs, nymphs, or adults). We identified 28 distinct genera in 20 families (Fig. 5, Table 1, Table S2), of which 26 are gram-negative, primarily represented by the Proteobacteria (16 genera distributed across the classes Alpha-, Beta-, and Gammaproteobacteria) and the Bacteriodetes (Fig. 6). Of the two gram-positive genera (phylum Firmicutes), Enterococcus was prevalent and abundant while Staphylococcus occurred at low levels (2-3%) in only some treatments (EYW and M; Table 1). Overall, five core genera are present in all sample types: Enterococcus, Chryseobacterium, Sphingobium, Delftia, and Sphingobacterium (Fig. 5A). These five accounted for one-third to fully two-thirds of the entire microbiome complement, depending on life history stage (Fig. 5B: central Venn diagram intersect). Consistent with patterns of species-level diversity (Fig. 4), egg-stage samples were comprised of up to twice as many bacterial genera as adult-stage samples, with 23 of the 28 genera present in eggs (untreated and/or surface-sterilized) compared to only 15 genera in adults (males and/or females, Fig. 5A,5C).

**Figure 5.**
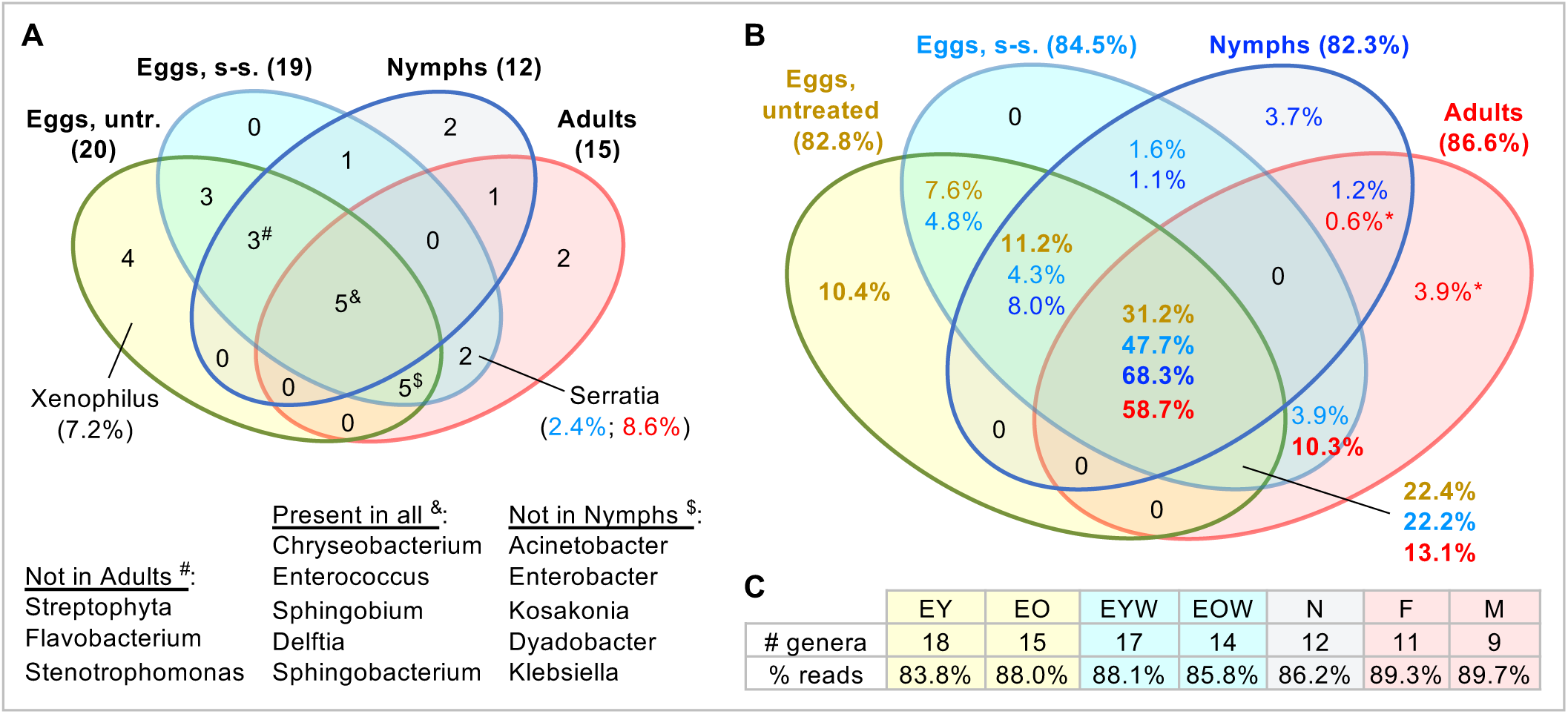
Distribution across life history samples of the 28 major genera of the *O. fasciatus* microbiome. Genera with >1% reads in at least one of the four major life history treatments are included. Venn diagrams are for **(A)** number of genera and **(B)** proportion of reads classified at the genus level. For diagram sectors with >10% reads, the genera are named, with selected abundances (% reads) specified parenthetically in (A). In (B), asterisks for low abundance in adult sectors reflect the result of averaging disparate abundance values between males and females (see also Fig. 7E, below). **(C)** For clarity, the genus counts and % reads are given for all seven individual stages and treatments. Note unclassified reads, as well as genera with <1% abundance, account for the approximately 15% of reads not documented here (*cf*., Fig. 3A,3C).

**Table 1.** The major prokaryotic genera detected across the life cycle in *O. fasciatus*. Twenty-eight genera were detected above the presence threshold of >1% abundance in at least one of the four major life history samples, and taxa are categorized by the number of life history samples in which they are present. For samples with <1% abundance, the exact value is specified but these values are shaded. Phylum abbreviations: Alpha/ Beta/ Gamma, Proteobacteria of the indicated class; Bact, Bacteriodetes; Cyano, Cyanobacteria; Firmi, Firmicutes.

| Taxonomy |  | Eggs, untr. |  | Eggs, s-s. |  | Nymphs | Adults |  | Distribution |
| --- | --- | --- | --- | --- | --- | --- | --- | --- | --- |
| Phylum | Genus | EY | EO | EYW | EOW | N | F | M |  |
| Bact | Chryseobacterium | 16.61% | 18.19% | 8.97% | 9.64% | 28.70% | 15.02% | 3.16% | Present in all |
| Alpha | Sphingobium | 7.13% | 2.42% | 1.23% | 2.26% | 15.18% | 14.61% | 1.18% |  |
| Firmi | Enterococcus | 3.38% | 3.40% | 28.08% | 29.62% | 20.15% | 28.56% | 49.47% |  |
| Beta | Delftia | 2.93% | 3.50% | 4.11% | 2.34% | 2.28% | 2.23% | 0.84% |  |
| Bact | Sphingobacterium | 1.90% | 2.90% | 3.33% | 5.91% | 1.96% | 1.33% | 0.97% |  |
| Gamma | Acinetobacter | 9.83% | 16.38% | 6.91% | 7.95% | 0.69% | 0.18% | 2.69% | Present in 3 (eggs and 1 post-embryonic stage) |
| Gamma | Enterobacter | 9.61% | 0.69% | 13.29% | 8.48% | 0.03% | 6.88% | 10.45% |  |
| Cyano | Streptophyta | 4.99% | 10.51% | 2.23% | 1.10% | 1.95% | 0.07% | 0.46% |  |
| Gamma | Kosakonia | 3.66% | 0.25% | 1.76% | 2.45% | 0.01% | 2.92% | 0.03% |  |
| Bact | Flavobacterium | 1.99% | 2.59% | 1.33% | 1.59% | 2.91% | 0.44% | 0.18% |  |
| Bact | Dyadobacter | 1.84% | 1.02% | 0.36% | 1.11% | 0.12% | 1.13% | 0.52% |  |
| Gamma | Stenotrophomonas | 1.75% | 0.63% | 1.46% | 0.87% | 3.18% | 0.18% | 0.73% |  |
| Gamma | Klebsiella | 1.42% | 0.10% | 0.85% | 1.13% | 0.00% | 1.17% | 0.19% |  |
| Other | Acidobacteria (Gp15) | 4.04% | 1.29% | 1.29% | 0.60% | 0.25% | 0.06% | 0.19% | Present in 2 |
| Gamma | Pseudomonas | 1.60% | 5.46% | 2.18% | 3.44% | 0.24% | 0.13% | 0.33% |  |
| Firmi | Staphylococcus | 0.93% | 0.51% | 2.09% | 0.95% | 0.81% | 0.03% | 3.27% |  |
| Alpha | Sphingomonas | 0.56% | 0.28% | 0.14% | 0.29% | 1.20% | 1.06% | 0.12% |  |
| Alpha | Brevundimonas | 0.41% | 2.42% | 1.28% | 0.77% | 0.83% | 0.16% | 0.07% |  |
| Alpha | Rhizobium | 0.13% | 0.23% | 2.44% | 0.81% | 1.11% | 0.35% | 0.35% |  |
| Gamma | Serratia | 0.12% | 0.50% | 2.78% | 1.94% | 0.00% | 11.50% | 5.76% |  |
| Beta | Xenophilus | 4.57% | 9.73% | 0.05% | 0.73% | 0.27% | 0.30% | 0.47% | Present in 1 |
| Beta | Comamonas | 1.37% | 0.89% | 0.43% | 0.47% | 0.19% | 0.21% | 0.32% |  |
| Bact | Epilithonimonas | 1.05% | 1.03% | 0.10% | 0.14% | 0.11% | 0.03% | 0.01% |  |
| Bact | Nubsella | 0.90% | 0.82% | 0.41% | 0.59% | 1.62% | 0.49% | 0.06% |  |
| Beta | Pseudorhodoferrax | 0.79% | 1.32% | 0.01% | 0.09% | 0.04% | 0.04% | 0.18% |  |
| Alpha | Shinella | 0.16% | 0.54% | 0.93% | 0.21% | 0.25% | 0.10% | 1.57% |  |
| Bact | Sediminibacterium | 0.12% | 0.00% | 0.00% | 0.00% | 0.08% | 0.01% | 6.19% |  |
| Other | Peredibacter | 0.05% | 0.45% | 0.04% | 0.30% | 2.12% | 0.09% | 0.01% |  |

**Figure 6.**
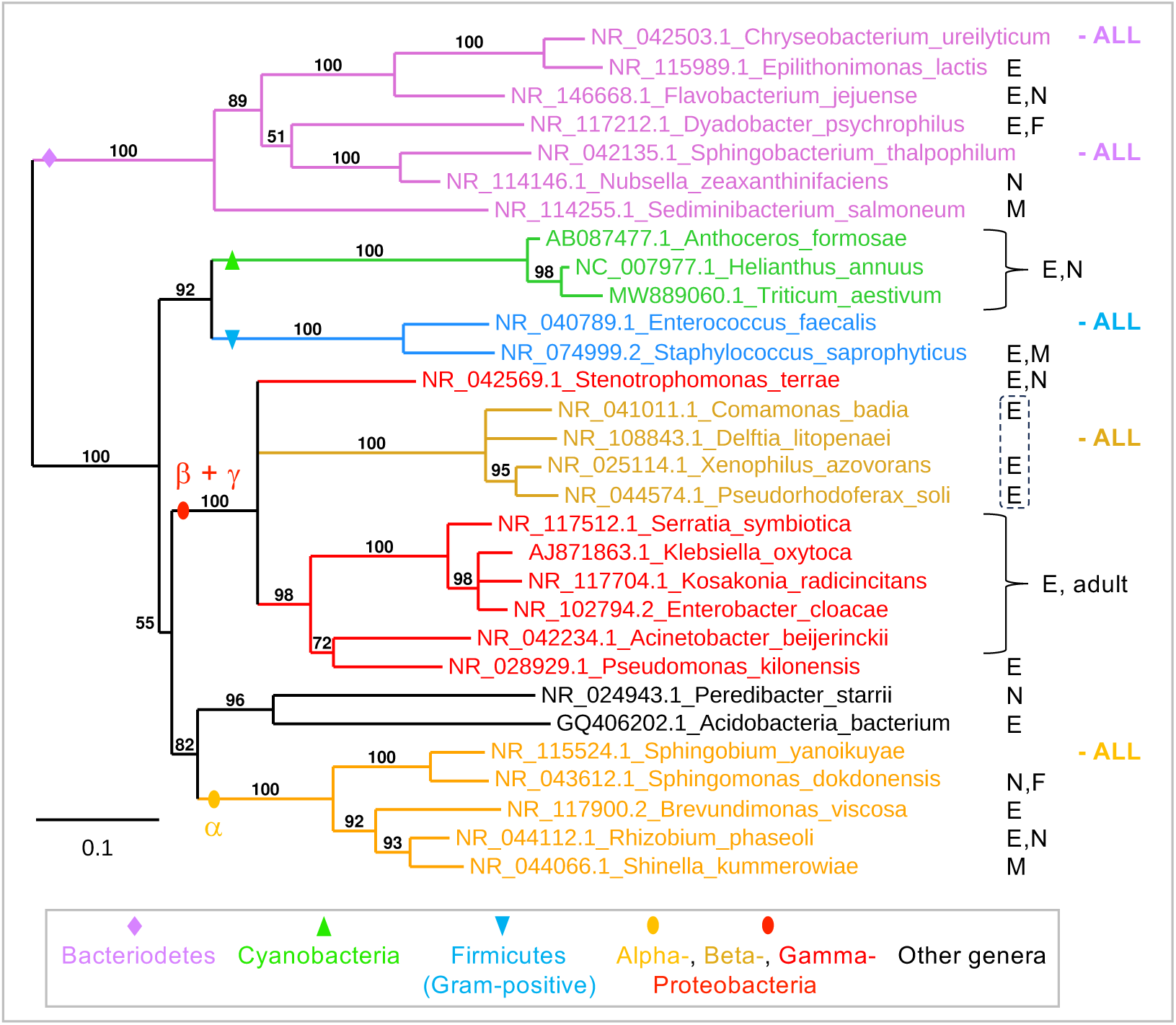
Phylogenetic distribution of the main microbiome genera. Maximum likelihood phylogeny for a >1350-bp region of the *16S rRNA* gene (species names and GenBank accessions as indicated); all nodes have ≥ 50% support; branch length unit is substitutions per site. As expected, bacterial taxa form well-supported clades at both the phylum and family levels. Taxa are annotated for presence across the *Oncopeltus* life cycle (stage: E, egg; N, nymph; F, female; M, male), as in Table 1. All genera are represented by a single species except the Cyanobacteria, where *Anthoceros formosae* is a molecular outgroup to the other two species (discussed in detail below).

To better assess microbiome diversity, we phylogenetically mapped the life history stage distribution of the 28 bacterial genera (Fig. 6). The five genera present throughout the milkweed bug life cycle are broadly distributed across the tree. The greater microbiome diversity at the egg stage chiefly arises from additional genera that are closely related to those main five. For example, all 11 genera in the clade of Beta- and Gammaproteobacteria are present at the egg stage. Except for Delftia being present throughout the life cycle, the other Betaproteobacteria – all specifically of the family Comamonadaceae – are restricted to embryonic samples (Fig. 6: dashed outline), and post-embryonic presence of any Gammaproteobacteria was patchy. Notably, no bacterial genera were uniquely present in either the surface-sterilized eggs or adult females (Fig. 5A, Table 1), so we next explored abundance dynamics of specific bacterial taxa to infer patterns of inheritance and acquisition.

### The egg surface and interior have distinct bacterial profiles

We examined egg-stage microbiome composition spatially and temporally. Greater relative abundance in untreated eggs reflects greater presence on the eggshell surface. Conversely, greater abundance in surface-sterilized samples is indicative of enrichment within the egg (yolk). We also assessed changes over time, as populations of bacterial taxa may expand or contract within the egg niche.

For both the untreated and surface-sterilized egg samples, five genera were lost and two other genera were gained in older eggs compared to young eggs (Fig. 5C, Table 1). Some of these changes likely reflect fluctuations relative to the 1% presence threshold, rather than composition turnover. However, genera with restricted presence (only in two life history stages, Table 1) may represent transient, opportunistic taxa in young egg samples (Acidobacteria in EY and EYW) or that proliferate on the eggshell surface (Brevundimonas in EO). Conversely, some genera are present in adult females and persist or thrive within eggs but are not maintained on the egg surface (Enterobacter, Kosakonia, and Klebsiella).

Stably over time, we find spatial differences in enrichment: within the egg for Serratia and Enterococcus; and on the egg surface for Chryseobacterium, Streptophyta, Comamonas, Epilithonimonas, and Xenophilus (Fig. 7A-B). The latter three genera are only present at the egg stage, and relative abundance of both Streptophyta and Xenophilus more than doubles on the egg surface during embryogenesis (Table 1). This suggests that, even in a simplified laboratory insect colony, the egg surface can foster taxa that are otherwise minimally present. On the other hand, both Enterococcus and Sphingobacterium are not only enriched within the egg (Fig. 7B) but also present throughout the milkweed bug life cycle (Fig. 5). This suggests that these taxa are transovarially transmitted as major taxa.

**Figure 7.**
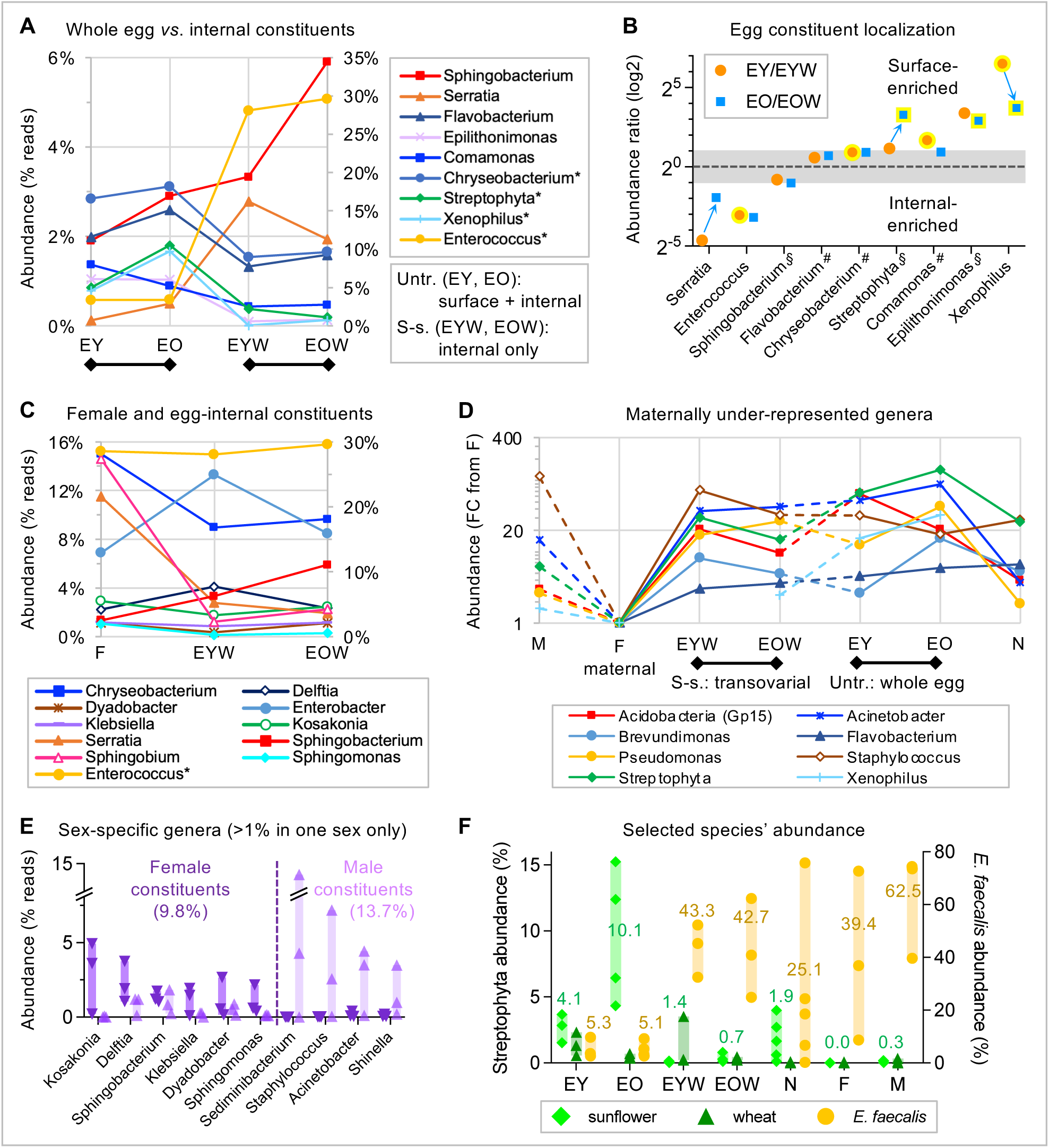
Life history stage-specific trends in bacterial abundance and transmission. **(A)** Mean abundance of egg-stage samples for genera that differ between untreated and surface-sterilized eggs: cool colors indicate greater abundance in the former, warm colors indicate greater abundance in the latter treatment (see legend, abbreviations as in Fig. 2). Genera with high abundance are denoted by an asterisk in the legend and plotted on the right-hand y-axis. Black lines below the plot indicate temporal progression within a treatment condition. **(B)** Locally enriched egg constituents are depicted as stage-specific ratios of untreated to surface-sterilized samples. Values outside the grey region have a >2-fold difference, which persists even in cases of changing relative abundance during embryogenesis (blue arrows). Yellow highlighting indicates significant differences (*p*<0.05) between untreated and surface-sterilized samples from unpaired, two-tailed comparisons; all tests were *t*-tests with Welch’s correction unless otherwise annotated: # for standard *t-*test (for EY/EYW and EO/EOW comparisons), § for non-parametric Mann-Whitney test (for EY/EYW comparisons only). **(C)** For the 11 genera present at >1% in adult females (F), mean relative abundance is shown in relation to transovarial transmission (EYW) and subsequent development (EOW). The genus with high abundance is denoted by an asterisk and plotted on the right-hand y-axis. **(D)** Eight genera are only detected at <0.5% in females yet are present at notable levels in eggs (>2% in ≥1 egg samples). Abundance is shown as fold change (FC) relative to adult female values. On the log10 scale, values <1 are not shown (for Brevundimonas and Flavobacterium in males or for Xenophilus in EYW and nymphs). Solid lines indicate developmental (temporal) continuity; dashed lines link samples that are not sequential. **(E)** Individual biological replicates (n=3) are plotted for the ten genera that differ in presence/absence between the sexes (adult females: dark purple, downward-pointing triangle plot points; adult males: light purple, upward-pointing triangles). Total sex-specific constituent values are the sum of the mean abundance per genus (as in Table 1). **(F)** Individual biological replicates (n=3-5) are plotted for each life history treatment, mapped to three selected species, as indicated in the legend. Note that the values here differ somewhat from those at the genus level (*e.g.,* other panels of this figure, Table 1), in part due to differences in read classification rates at different taxonomic levels (Fig. 3A). Values are mean species abundance for *E. faecalis* (yellow text) or the sum of mean abundance values for both sunflower and wheat as Streptophyta constituents (green text).

### The egg stage microbiome does not simply recapitulate the maternal profile

Regarding potential transovarial transmission, of the eleven genera present in females (Table 1), ten are also detected within surface-sterilized eggs (Fig. 7C, the exception being Sphingomonas). However, over developmental time (across F, EYW, EOW), relative abundance is variously: stable (Enterococcus and Kosakonia), fluctuates in young eggs before returning to maternal levels in late embryogenesis (greater in EYW: Delftia and Enterobacter, lower in EYW: Klebsiella and Dyadobacter), less abundant in eggs than females (Chryseobacterium, Sphingobium, Serratia), or increases (Sphingobacterium). Thus, variable abundance dynamics argue against transmission of a specific bacterial community as a suite of correlated taxa at the maternal to embryonic life history stage transition.

Furthermore, as noted above, female bacterial taxa only represent a fraction of egg stage bacterial diversity (Figs. 4, 5). We thus evaluated taxon-specific abundance across all life history samples for eight genera with very low female abundance (<0.5%) and >4× greater egg stage abundance (>2% in at least one egg-stage sample). Relative to adult females, all eight genera are more abundant at other life history stages (Fig. 7D), but only half of these have >1% abundance in either a nymphal or adult male sample. This suggests that some egg-stage bacterial genera are unique – neither enriched in the female reproductive tract [33, 36] nor in other postembryonic bacteria-harboring tissues [54].

### Sex-specific differences in the adult microbiome profile

Adults have the lowest bacterial diversity of all life history stages (Fig. 4), with five shared bacterial genera accounting for the majority of the microbiome (77% in females, 70% in males; Table 1). However, fully 10-14% of the adult microbiome is sex-specific, represented by ten genera (Fig. 7E). Most are variably present, with below-threshold abundance (<1%) in at least one biological replicate. Only Delftia and Sphingobacterium were consistently present (>1%) in all female replicates – as well as in at least one male replicate, supporting their inclusion as part of the core microbiome (Fig. 5A). Despite variability between individuals, we do detect striking sex-specific differences. Compared to a scarcely-detectable <0.03% in the other sex, individual males harbored up to 7% Staphylococcus or 14% Sediminibacterium, with Kosakonia as the major female-specific constituent at up to 5%.

### Species-level prokaryotic profiles highlight taxon proportionality and environmental components

Certain additional patterns in the *O. fasciatus* microbiome emerge in species-level analyses, with bacterial abundance differing across compared to within genera. Chryseobacterium is present in all life history samples (Table 1), but with high variation (range: 0.83-56.3% in post-embryonic individuals, n=9, Supplementary Data File S1). Nonetheless, species’ proportions are remarkably uniform. Consistently, the two most abundant species account for 94% of Chryseobacterium reads: *C. tructae* at 50.7 ± 1.5% and *C. hominis* at 43.4 ± 1.9% (median ± median absolute deviation, among 26 total Chryseobacterium species, Supplementary Data File S1). Thus, even when a genus varies as a fraction of the total microbiome, the proportions of major species are consistent within the genus.

This pattern is corroborated by the major constituent Enterococcus (Figs. 3C and 7A-C, Table 1). *E. faecalis* is the dominant species (of 13), with >97% of Enterococcus reads (Supplementary Data File S1). Nonetheless, *E. faecalis* abundance is extremely variable (Fig. 7F). In individual post-embryonic insects (n=9), *E. faecalis* abundance ranged from an astonishing >72% of the microbiome (four individuals: one nymph, one adult female, two adult males), to <10% (two individuals), to only 0.2% in one nymph. Given our experimental design of simultaneous sampling for all individuals of a given life history stage, these are striking differences in microbiome composition at the intra-specific, within-population level.

Microbiome evaluation from a uniformly simple lab colony also reveals stage-specific importance of environmental components. We detect plant chloroplast *16S rRNA* (phylum Cyanobacteria, genus Streptophyta: Figs. 3B, 6). This is a known issue in microbiome profiling of phytophagous arthropods, including seed-feeding insects, and is generally regarded as contamination [55]. Yet, whole body or egg clutch material is a form of holobiont or environmental sample. In all five non-adult samples (EY, EO, EYW, EOW, N), Streptophyta accounted for 1-11% of genus-level reads (Table 1, Fig. 7F). This contrasts with negligible detection in any adult sample (<0.34% in five of six samples; Fig. 7F, Supplementary Data File S1), despite uniform handling of nymphal and adult material (see Methods). Moreover, Streptophyta is a defining constituent of untreated eggs, doubling in abundance during embryogenesis, from 5.0% to 10.5% (Fig. 7A-B, Table 1). At the species level, sunflower (*Helianthus annuus*) and wheat (*Triticum aestivum*) accounted for up to 95% of Streptophyta reads in all samples (Supplementary Data File S1). Low-level detection of wheat likely does represent minor contamination from milled flour dust, from food stocks for other insect species maintained in the same lab rooms. In contrast, sunflower seeds are the food source for the milkweed bug colony and thus a direct environmental component. As detection of sunflower *16S* material was stage-specific but otherwise very low (see next section), we consider this to be a specific environmental feature (see Discussion).

### Representative sampling with minimal sequencing, including for environmental material

Finally, we used Sanger sequencing and GenBank blastn as an independent analysis and classification approach, clarifying environmental levels of Streptophyta.

In insect samples we detected a good balance of major, abundant taxa and rare, but genuine, constituents (Table S1). For example, Sanger sequencing of nymphal sample N11 recovered Enterococcus (66.2% in the full-scale dataset) as well as Sphingobium (second-most abundant nymphal genus) and Enterobacter (rare nymphal genus). Across life history samples, 97% of clones belonged to the 28 main genera (Table 1), with 86% to the ten most abundant. One clone was classified outside the main genera but was still widely detected (Acidovorax: 0.1-0.6% abundance, 380-2000 reads per sample). For surface-sterilized eggs, either technical replicate detected multiple clones of a highly abundant genus and an additional genus with >1% abundance (Tables 1, S1). Overall, the 37 sequenced clones identified ten distinct bacterial genera. Thus, while small-scale sequencing clearly will not document the full scope of the microbiome, it was more effective than anticipated (*cf.*, [33, 56]) at detecting a meaningful variety of taxa.

Building on this framework, we sequenced samples taken directly from laboratory food stocks for our insect cultures, obtaining good sensitivity and accuracy. All nine clones from two biological replicates of sunflower seeds exclusively identified *Helianthus annuus* chloroplast DNA, while all five clones from a sample of milled flour only identified chloroplasts of the grass family Poaceae (a mix of wheat and maize, Table S1).

Notably, we did not detect any chloroplast (Streptophyta) material in Sanger sequencing of the insect-derived samples, which included the two biological replicates with the highest levels of Streptophyta in the full-scale dataset (samples EO6 and EO7, with 17.0% and 12.8% Streptophyta, respectively; Supplementary Data File S1). We therefore conclude that stage-specific detection of sunflower material is a specific attribute of these samples, rather than detection of general environmental material.

## DISCUSSION

We present a nuanced account of the microbiome of the milkweed bug *O. fasciatus*, a Lygaeinae species that lacks both midgut crypts and bacteriocytes. In characterizing the core microbiome profile, we assess the implications of changing abundance between individuals and across life history stages for modes of bacterial acquisition.

### Microbiome repertoires are dynamic across the life cycle, with complex acquisition

By assessing the microbiome profile at each of the phylum, family, genus, and species levels (Figs. 3-7), we dissected several dynamic trends, including high bacterial diversity at the egg stage that declines postembryonically. We also find notable differences across the life cycle for bacterial repertoires: within or external to the egg, between the sexes, and for stage-specific levels of variation between individuals (Figs. 3B-C, 7). Thus, attention to sampling and numerical handling are important for identifying patterns of bacterial abundance from noisy data, even for biological replicates derived from controlled lab colony environments.

Is vertical transmission predominantly transovarial, directly into the egg, or via maternal secretions onto the eggshell surface? We find conflicting evidence for the maternal bacterial repertoire. The five most abundant genera in females (each >6%) have mixed localization profiles in eggs, with three internally enriched and two externally enriched. Of the six other genera in females, four are externally enriched in young eggs – consistent with egg-smearing transmission. However, these taxa decline externally and increase internally during embryogenesis, supporting the importance of transovarial transmission. Equally, egg-stage bacterial genera exhibit conflicting patterns. Internally-enriched taxa include Enterococcus and Serratia, which are characteristic of both males and females. As shared bacterial taxa in adults are few (Fig. 7E), and this is a persistent egg-stage feature, these could be key constituents. However, absence of Serratia in nymphs challenges this. Aside from Chryseobacterium as a stable constituent across the life cycle, no other egg surface-enriched bacteria are present (>1%) in females, suggesting they are environmental constituents of the egg niche or that these taxa are vastly outcompeted within the female. Interestingly, in the pea aphid the obligate endosymbiont Buchnera and the facultative endosymbiont Serratia (also detected in *O. fasciatus*) differ in their mode of transmission from the female to the egg [32]. Hence the diversity of transmission routes seen across the Hemiptera [30, 31] may also pertain even within a single species, belying the expectation for a global pattern.

To what extent is bacterial relative abundance important? In some cases, physiologically important bacteria are prevalent, such as Burkholderia symbionts in the fellow bug *Riptortus pedestris* [56]. Indeed, abundance can be a direct method of benefit to the insect host (“colonization resistance”), such as through chemically dictating the environment (*e.g.*, acidic or alkaline, [reviewed in 57]), or by outcompeting potentially pathogenic bacteria, as seen for *E. faecalis* in the silk moth *Bombyx mori* [reviewed in 45]).

However, there need not be a correlation between abundance and biological relevance. The high levels – and large inter-individual fluctuations – that we report here in constituents like *E. faecalis* argue for targeted approaches. This could include filtering highly abundant taxa. For example, when excluding Enterococcus from our dataset, 75% of biological replicates (18 of 24) gained bacterial genera at >1% abundance (1-11 additional genera per sample). Bacterial diversity more than doubled for the individual nymphal and male samples that were dominated by Enterococcus (>56%). Across all samples, half of the gains were relative increases that reinforce the prevalence of the core 28 genera (Table 1). Among additional genera gained after filtering, all post-embryonic individuals with >56% Enterococcus showed an increase in the fellow gram-positive, lactic acid bacteria Lactobacillus and Vagococcus. Gains in Proteobacteria included Pseudoxanthomonas in nymphs (closely related to the nymph-enriched core genus Stenotrophomonas). Intriguingly, after filtering we also detect Polaromonas, another member of the egg-characteristic Comamonadaceae family, in both F and EYW samples, suggesting this too may be a potential transovarially transmitted constituent. Thus, filtering may reveal meaningful bacterial diversity that is otherwise concealed by the most abundant genera.

### Detection of chloroplast material: more than simple contamination

We report consistent, notable detection of chloroplast *16S* DNA from the insect’s sunflower seed food source in untreated eggs and nymphs, but not in surface-sterilized eggs or adults (Fig 7F). Neither the spatial structure of the insect colonies (oviposition into cotton wool, remote from the food dish) nor the dormant state of the seeds (neither germinating nor dusty) accounts for specific egg-stage surface or nymphal detection. Clearly, the eukaryotic food source is not a component of the microbiome. Nonetheless, we emphasize this *16S* detection pattern as a potentially defining characteristic for how nutritional material may be sequestered (by nymphs?) or provisioned (by females at the oviposition site?) in a life history stage specific manner (and see the next section for speculation on its potential relevance).

### Comparisons with other insects’ microbiomes: shared genera and nitrogen-fixing taxa

We compared the core milkweed bug microbiome with reports of the predominant genera in other insects. Overall, *O. fasciatus* has a fairly typical insect and hemipteran microbiome profile, albeit without well-known symbionts of fellow hemipterans. Surprisingly, this is in part irrespective of feeding ecology type.

With the closely related firebug *Pyrrhocoris apterus*, 16 bacterial genera are shared (Table S2). This nuanced study on the firebug compared bacterial profiles across different geographical sites, life history stages, segments of the midgut, and lab colony diets, as well as environmental sampling of the primary linden seed food source [58]. As in our findings in the milkweed bug, the highest levels of bacterial diversity in the firebug occurred at the egg stage, where “other” genera outside of the most prevalent 15 accounted for fully a third of the microbiome profile at this stage. Furthermore, when *P. apterus* was also raised on a diet of sunflower seeds, there was a greater abundance of Acinetobacter, Delftia, and Sphingobium, all of which are major constituents in *O. fasciatus*. In environmental analysis of the natural linden seed food source, bacteria on the seeds included several prominent constituents in the *P. apterus* midgut. Regardless of whether this represents undigested food or part of the firebug microbiome, five of these genera are also present in the milkweed bug. In sum, a large proportion of the milkweed bug core microbiome is shared with a fellow seed-feeding bug, and a number of these bacterial taxa are closely associated with the diet.

More widely, we found a high level of microbiome commonality not only with other hemipterans but also with a carnivorous hemimetabolous insect as well as phytophagous holometabolous species (Table S2, 71 species in five insect orders, [50, 54, 57–63]). Pseudomonas and Acinetobacter occur widely across these diverse insects. While prevalent, Enterococcus is neither ubiquitous nor necessarily represented by *E. faecalis* (*E. mundtii* dominates in the moth *Spodoptera littoralis*, [50]). Several taxa were not present in the carnivorous earwig (Dermaptera, [62]) but are frequently detected across the Hemiptera and Holometabola (selected Coleoptera, Lepidoptera, Diptera), including Serratia, Enterobacter, Klebsiella, and Sphingomonas. Yet, eight genera are shared between the milkweed bug and earwig. This contrasts with limited overlap (only three genera) with either a rice-feeding leafhopper [54] or a seed-feeding stinkbug [60] as fellow hemipterans.

Interestingly, *O. fasciatus* harbors two nitrogen-fixing, soil-associated bacterial genera: Klebsiella [64, 65], widely occurring in insects, and Rhizobium, which may be a specific hemipteran (seed-feeding bug) constituent (Tables 1 and S2, [58, 60, 61]). In previous work on insect’s capacities for nitrogen metabolism, we observed that *O. fasciatus* and other hemipterans’ genomes do not encode the metabolic enzymes for synthesis of arginine, and we suggested that this was obviated by consuming a nutrient-rich seed diet [11, 66]. However, it is tempting to now speculate that particularly Rhizobium may indirectly derive nutritional benefits from association with seed-feeding insects, which could be consistent with our observations on structured detection of chloroplast material. In turn, Rhizobium may provide these Hemiptera with nitrogen metabolism components that they lack, providing nutritional symbiosis and complementation.

In sum, this selected comparison documents 20 of the 27 milkweed bug core bacterial genera in other insects (Table S2). Five other genera were detected with low abundance in few samples. Exceptions to this are Kosakonia, which is prevalent across milkweed bug life history stages, and Xenophilus, which is notably abundant in untreated eggs (3.5-12.7% per sample) as part of the greater Comamonadaceae diversity revealed in our focused egg-stage microbiome sampling. Future, targeted research will help to clarify which of these bacterial taxa comprise genuine nutritional symbionts (obligate or facultative), and which are merely commensal or even pathogenic [45, 57].

Finally, we considered known hemipteran nutritional endosymbionts. However, these bacterial genera were not detected or negligible (as low as <0.008%). Specifically, we do not detect: Burkholderia, environmentally acquired in many stinkbugs [34, 67, 68]; the kissing bug *Rhodnius prolixus*’s Rhodococcus [69]; or Candidatus endosymbionts, known in several lygaied and stinkbugs [29, 39]. Unsurprisingly, we do not detect the pea aphid *Acyrthosiphon pisum*’s exclusive, obligate symbiont Buchnera [20]. Wolbachia, which provides the bed bug *Cimex lectularius* with nutritional benefits aside from its well-known, widespread influence on insect sex determination [70], is also not present in *O. fasciatus*.

### Conclusions

Our controlled sampling of the microbiome from a lab colony of the milkweed bug documents high variability between individual insects for some bacterial taxa – a feature that has been understudied to date [discussed in 45]. We also find marked differences in the overall microbiome repertoire and diversity between life history stages, even though as a hemimetabolous species without complete metamorphosis *O. fasciatus* exhibits no major changes in anatomy, behavior, or ecology across the life cycle. Nonetheless, we define a robust core of 28 prokaryotic genera in *O. fasciatus*, most of which are typical for insects. As in previous work [e.g., 38, 58], direct visualization of bacteria within *O. fasciatus* as well as tissue-specific *16S* profiling will clarify where bacteria are housed and the extent to which selected taxa may be transovarially transmitted. Also of note is the prevalence of nitrogen-fixing Rhizobium in several Hemiptera, and in *O. fasciatus* the prevalence and diversity of the Comamonadaceae family at the egg stage. It will be interesting the examine the extent to which these trends are borne out for *O. fasciatus* feeding on its native food source of cardenolide-containing *Asclepias* wildflower seeds in natural populations.

## SUPPLEMENTARY MATERIAL

### Supplementary PDF file

**Table S1.**
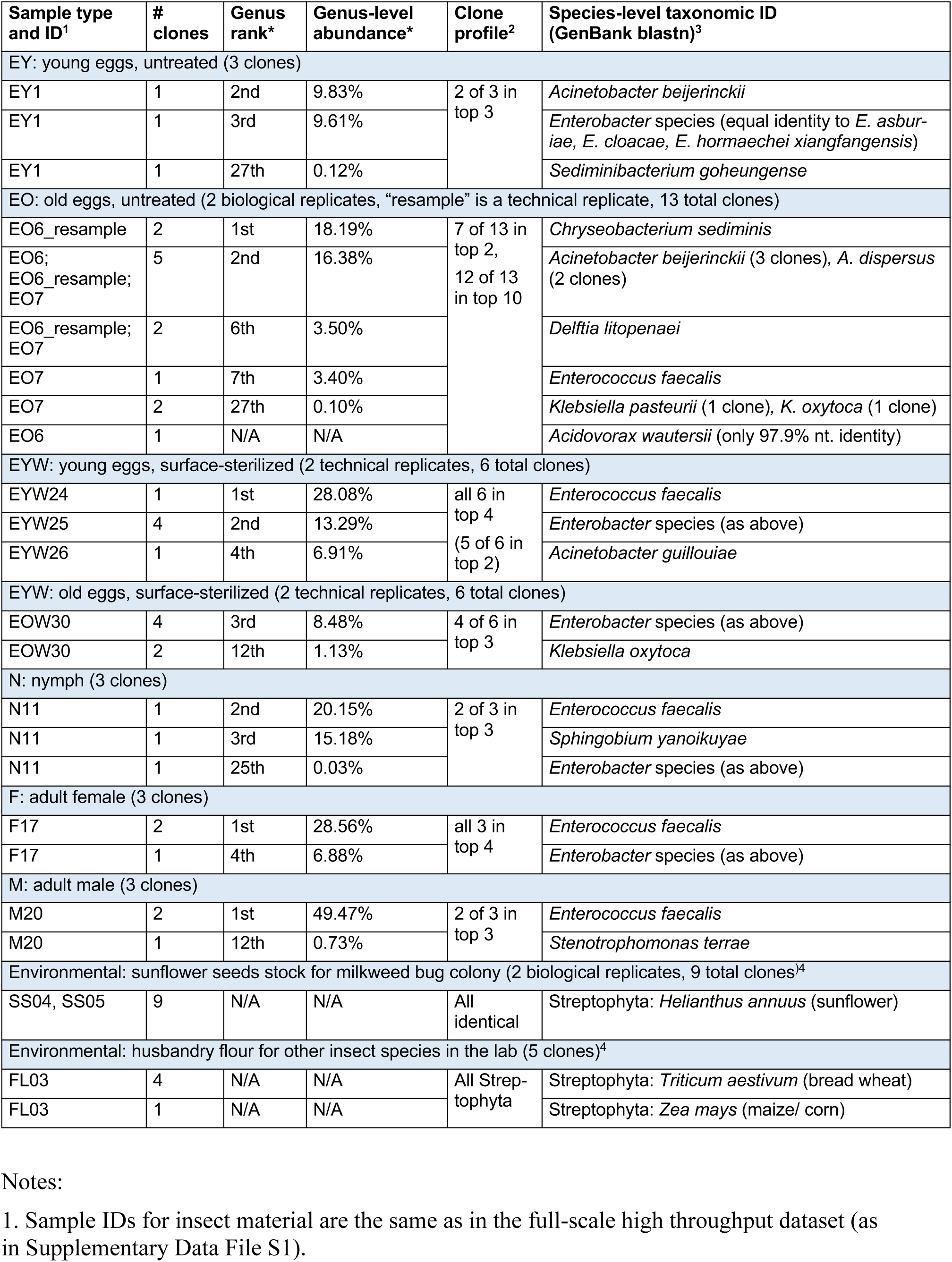
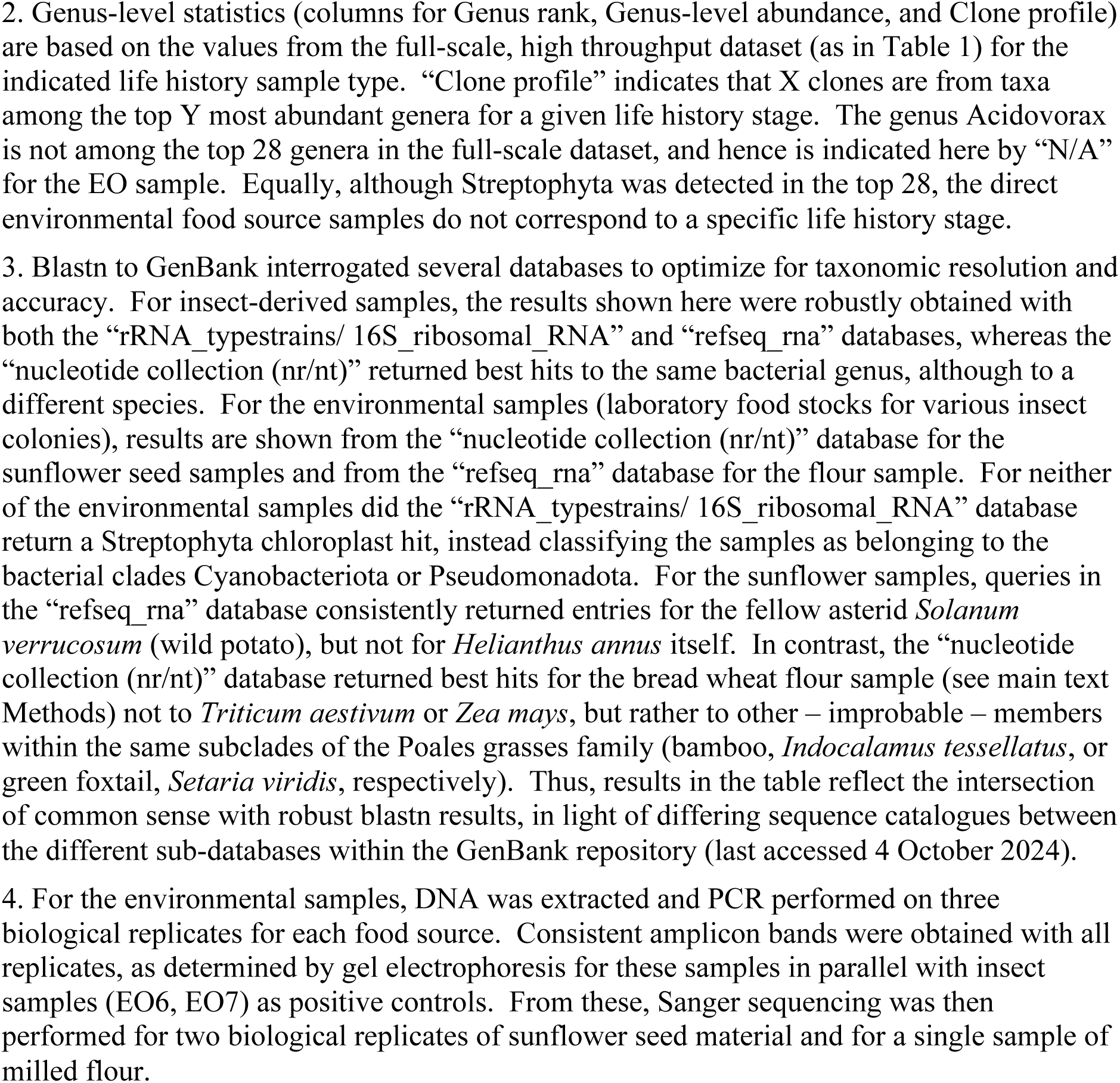
Sanger-sequencing results. Biological and technical replicates as indicated per life history stage or environmental sample type, with 3-5 clones per replicate.

**Table S2.**
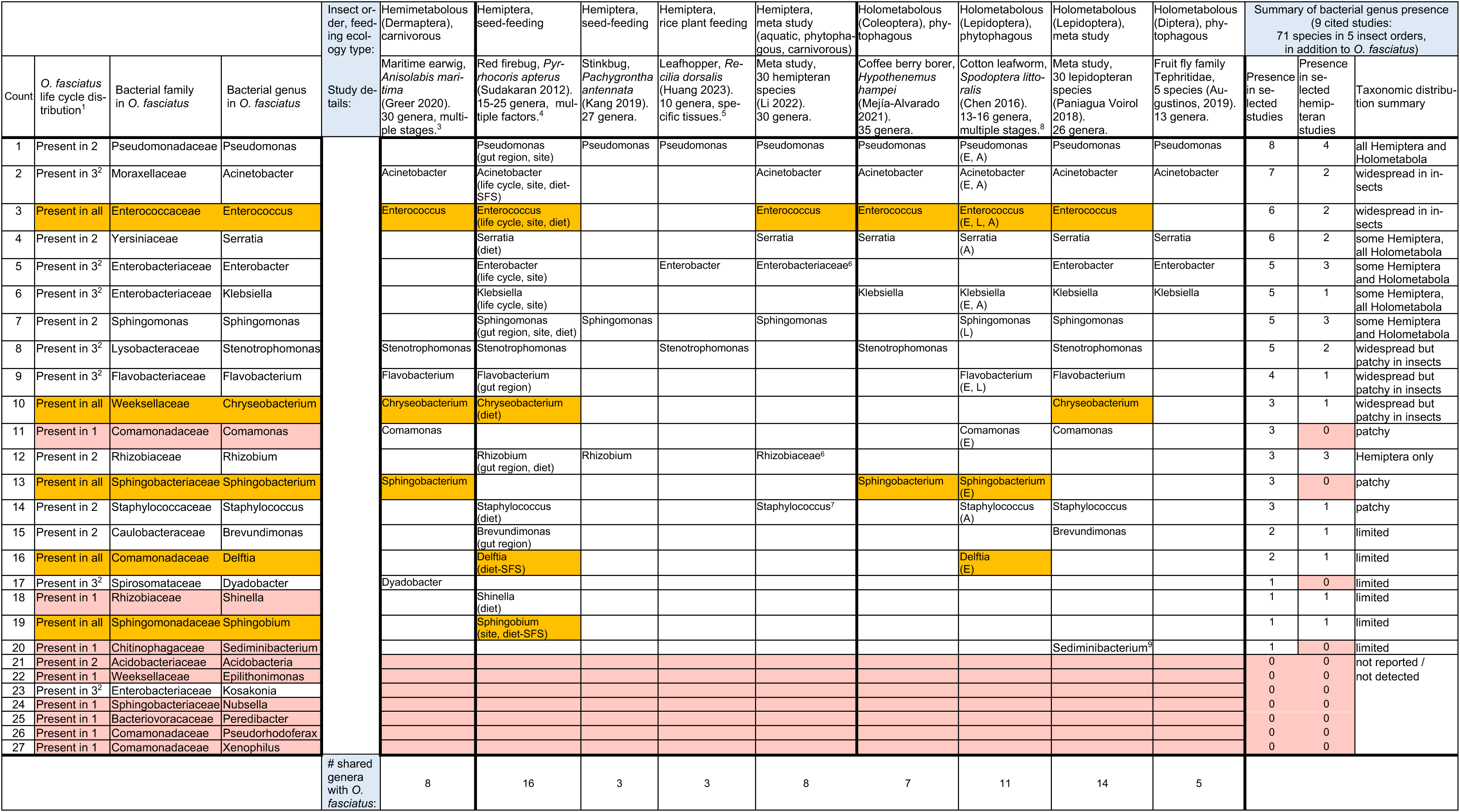
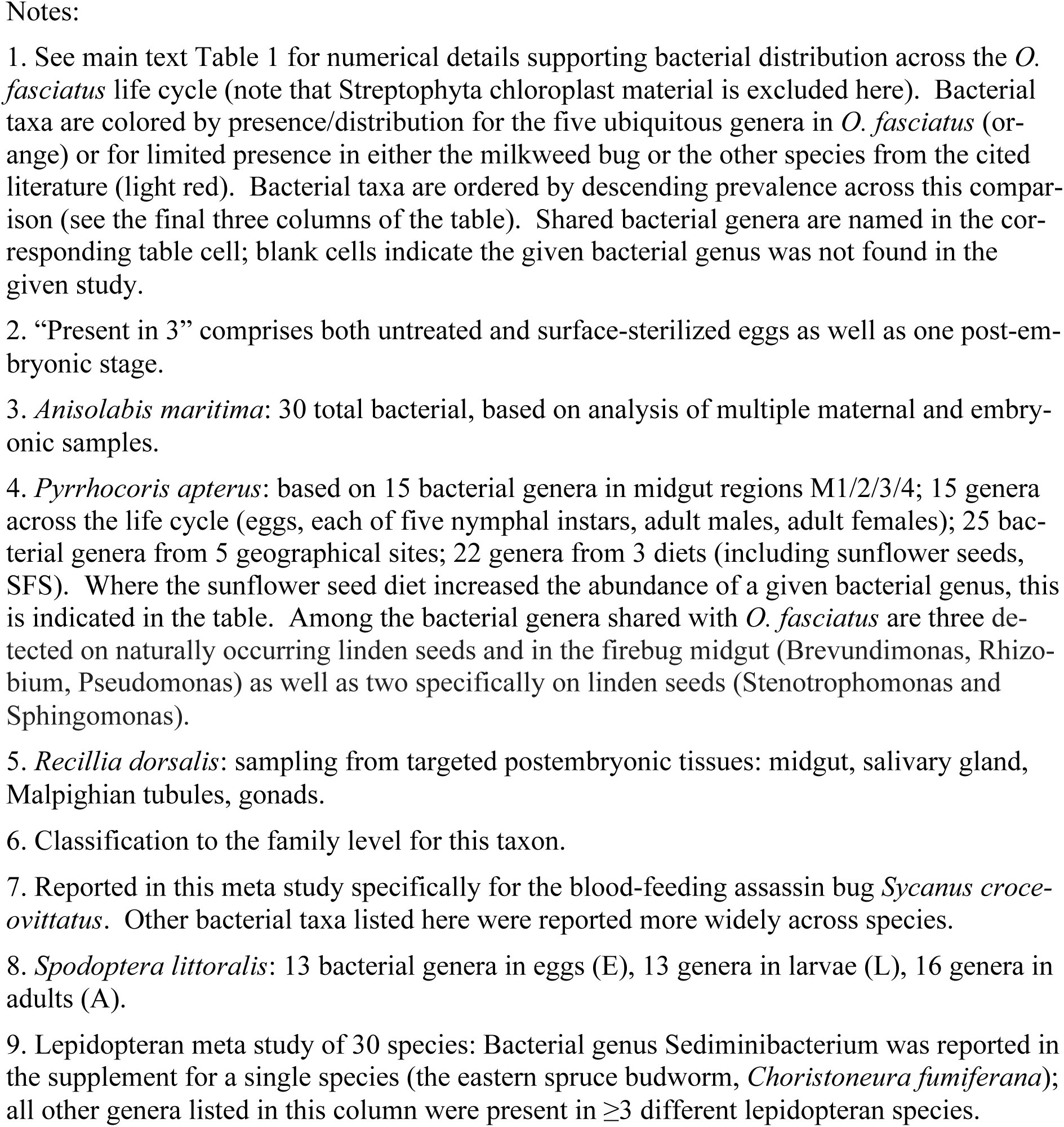
Comparison of bacterial genera shared between *O. fasciatus* and selected insect species. Bacterial profiles are based on the major bacterial genera reported in main text display elements of the cited studies, unless otherwise noted [1–9]. Streptophyta (chloroplast) material is not considered here.

### Supplementary Excel spreadsheet

**Supplementary Data File S1. Primary data on *16S rRNA* read classification.**

## STATEMENTS and DECLARATIONS

## Acknowledgments

We thank: Poornima Roy for extremely helpful suggestions on experimental design and discussions on data interpretation throughout the project; Christian Becker of the Cologne Center for Genomics for overseeing library preparation and sequencing, discussions on MiSeq data quality, and data access support; and Georg Petschenka and Amelia Silva for insightful discussions on the biological interpretation of microbiome profiles.

## Data availability

The data generated or analyzed during this study are included in the manuscript (and its supplementary information files).

## Author contributions

WL developed sample preparation methodology, generated and analyzed the data, and contributed to primary writing of the paper and editing of the paper. NTdSG conducted orthology searches and phylogenetic analyses and contributed to editing of the paper. KAP conceived and supervised the study, developed sample preparation methodology, analyzed the data, and contributed to primary writing of the paper and editing of the paper.

## Supplementary Information

In this PDF:

### Additional supplementary file

Supplementary Data File S1. Primary data on *16S rRNA* read classification (Excel file).

All sequenced samples are identified by life history sample type (EY, EO, EYW, EOW, Ni, Np, N all, F, M, as in main text Fig. 2), as well as by technical replicate (K or R, depending on polymerase, as described in the Methods), and with a unique number ID per sample.

Numerical details on sequencing yields and classification are provided in four tabs:

- <u>Read depth statistics</u>: Reports the total number of classified reads at the species level as well as the associated Shannon Species Diversity Index and total number of species identified, for all 48 samples and evaluated per technical replicate. Descriptive statistics as well as calculations and plots of correlation coefficients and t-test comparisons between technical replicates are provided. As we find no correlation between read depth and species diversity, we consider that sufficient read depth was obtained for all samples.
- <u>Phylum (%, counts)</u>: Organized from left to right, the number of reads assigned to each phylum is presented for: the mean % abundance across biological replicates, the mean % abundance across technical replicates, the mean % abundance per sample, and finally the read counts per sample. The 6 most abundant phyla are highlighted in light yellow. Grey shading indicates abundance <1% (for mean % abundance, columns B:J) or 0 reads (for read counts per sample, columns CE:DZ).
- <u>Genus (%, counts)</u>: Organized from left to right in the same manner as for the Phylum tab. The main 28 genera are highlighted in light yellow. Grey shading indicates abundance <1% (for mean % abundance, columns B:J). Red and green shading indicate <10 reads or >1000 reads, respectively (for read counts per sample, columns CE:DZ).
- <u>Species (counts)</u>: Read counts per sample and per species are presented. Species are listed alphabetically, with highlighting for genera and species featured in the main text: Chryseobacterium (light yellow) and *C. tructae* and *C. hominis* (dark yellow), Enterococcus (light orange) and *E. faecalis* (bright orange), Streptophyta (light green) and *Helianthus annuus* and *Triticum aestivum* (bright green). Grey shading indicates 0 reads.

